# Neanderthal-Denisovan ancestors interbred with a distantly-related hominin

**DOI:** 10.1101/657247

**Authors:** Alan R. Rogers, Nathan S. Harris, Alan A. Achenbach

## Abstract

Previous research has shown that modern Eurasians interbred with their Neanderthal and Denisovan predecessors. We show here that hundreds of thousands of years earlier, the ancestors of Neanderthals and Denisovans interbred with their own Eurasian predecessors—members of a “superarchaic” population that separated from other humans about 2 mya. The superarchaic population was large, with an effective size between 20 and 50 thousand individuals. We confirm previous findings that: (1) Denisovans also interbred with superarchaics, (2) Neanderthals and Denisovans separated early in the middle Pleistocene, (3) their ancestors endured a bottleneck of population size, and (4) the Neanderthal population was large at first but then declined in size. We provide qualified support for the view that (5) Neanderthals interbred with the ancestors of modern humans.

**One-sentence summary:** We document the earliest known interbreeding between ancient human populations and an expansion out of Africa early in the middle Pleistocene.

## Introduction

During the past decade, we have learned about interbreeding among hominin populations after 50 kya, when modern humans expanded into Eurasia [1, 2, 3]. Here, we focus farther back in time, on events that occurred more than a half million years ago. In this earlier time period, the ancestors of modern humans separated from those of Neanderthals and Denisovans. Somewhat later, Neanderthals and Denisovans separated from each other. The paleontology and archeology of this period record important changes, as large-brained hominins appear in Europe and Asia, and Acheulean tools appear in Europe [4, 5]. It is not clear, however, how these large-brained hominins relate to other populations of archaic or modern humans [6, 7, 8, 9]. We studied this period using genetic data from modern Africans and Europeans, and from two archaic populations, Neanderthals and Denisovans.

Fig. 1 illustrates our notation. Upper-case letters refer to populations, and combinations such as *XY* refer to the population ancestral to *X* and *Y*. *X* represents an African population (the Yorubans), *Y* a European population, *N* Neanderthals, and *D* Denisovans. *S* is an unsampled “superarchaic” population that is distantly related to other humans. Lower-case letters at the bottom of Fig. 1 label “nucleotide site patterns.” A nucleotide site exhibits site pattern *xyn* if random nucleotides from populations *X, Y*, and *N* carry the derived allele, but those sampled from other populations are ancestral. Site pattern probabilities can be calculated from models of population history, and their frequencies can be estimated from data. Our Legofit [10] software estimates parameters by fitting models to these relative frequencies.

**Figure 1:**
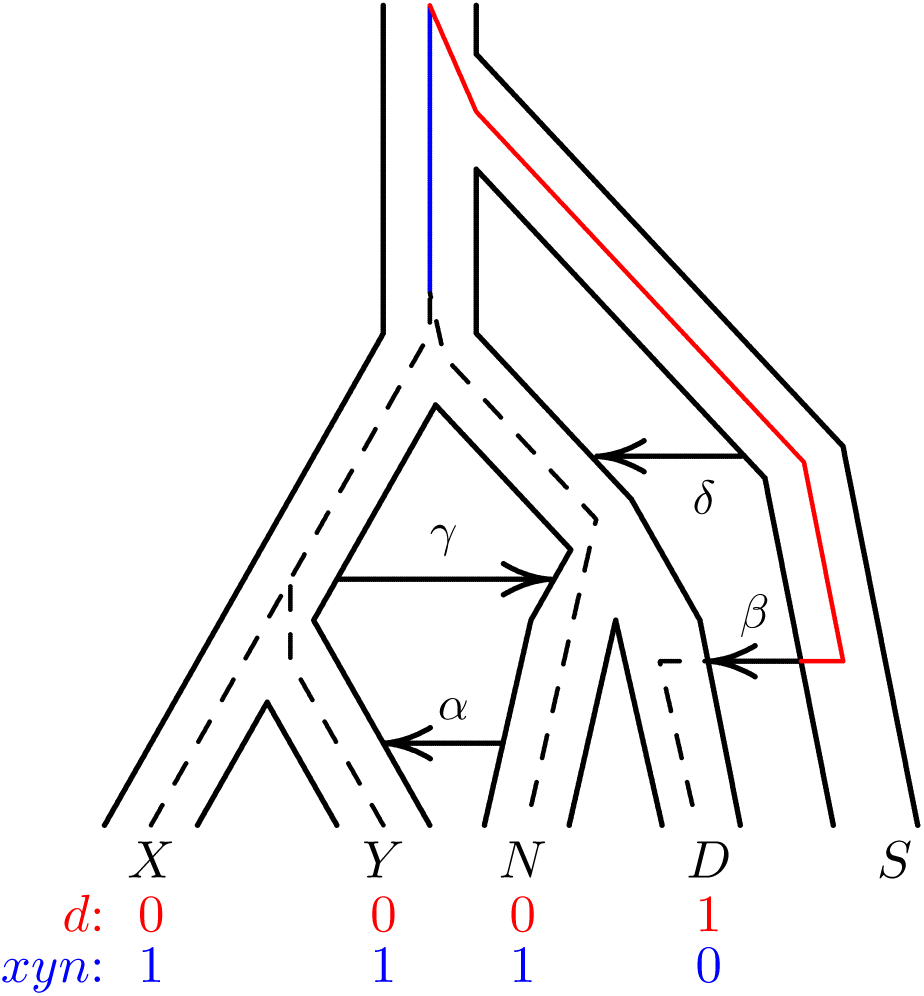
A population network including four episodes of gene flow, with an embedded gene genealogy. Upper case letters (*X, Y, N, D*, and *S*) represent populations (Africa, Europe, Neanderthal, Denisovan, and superarchaic). Greek letters label episodes of admixture. *d* and *xyn* illustrate two nucleotide site patterns, in which 0 and 1 represent the ancestral and derived alleles. A mutation on the red branch would generate site pattern *d*. One on the blue branch would generate *xyn*. For simplicity, this figure refers to Neanderthals with a single letter. Elsewhere, we use two letters to distinguish between the Altai and Vindija Neanderthals.

Nucleotide site patterns contain only a portion of the information available in genome sequence data. This portion, however, is of particular relevance to the study of deep population history. Site pattern frequencies are unaffected by recent population history, because they ignore the within-population component of variation [10]. This reduces the number of parameters we must estimate and allows us to focus on the distant past.

The current data include two high-coverage Neanderthal genomes: one from the Altai Mountains of Siberia and the other from Vindija Cave in Croatia [13]. Rather than assigning the two Neanderthal fossils to separate populations, our model assumes that they inhabited the same population at different times. This implies that our estimates of Neanderthal population size will refer to the Neanderthal metapopulation rather than to any individual subpopulation.

The Altai and Vindija Neanderthals appear in site pattern labels as “*a*” and “*v*”. Thus, *av* is the site pattern in which the derived allele appears only in nucleotides sampled from the two Neanderthal genomes. Fig. 2 shows the site pattern frequencies studied here. In contrast to our previous analysis [14], the current analysis includes singleton site patterns, *x, y, v, a*, and *d*, as advocated by Mafessoni and Prüfer [15]. A simpler tabulation, which excludes the Vindija genome, is included as SOM Fig. S2.

**Figure 2:**
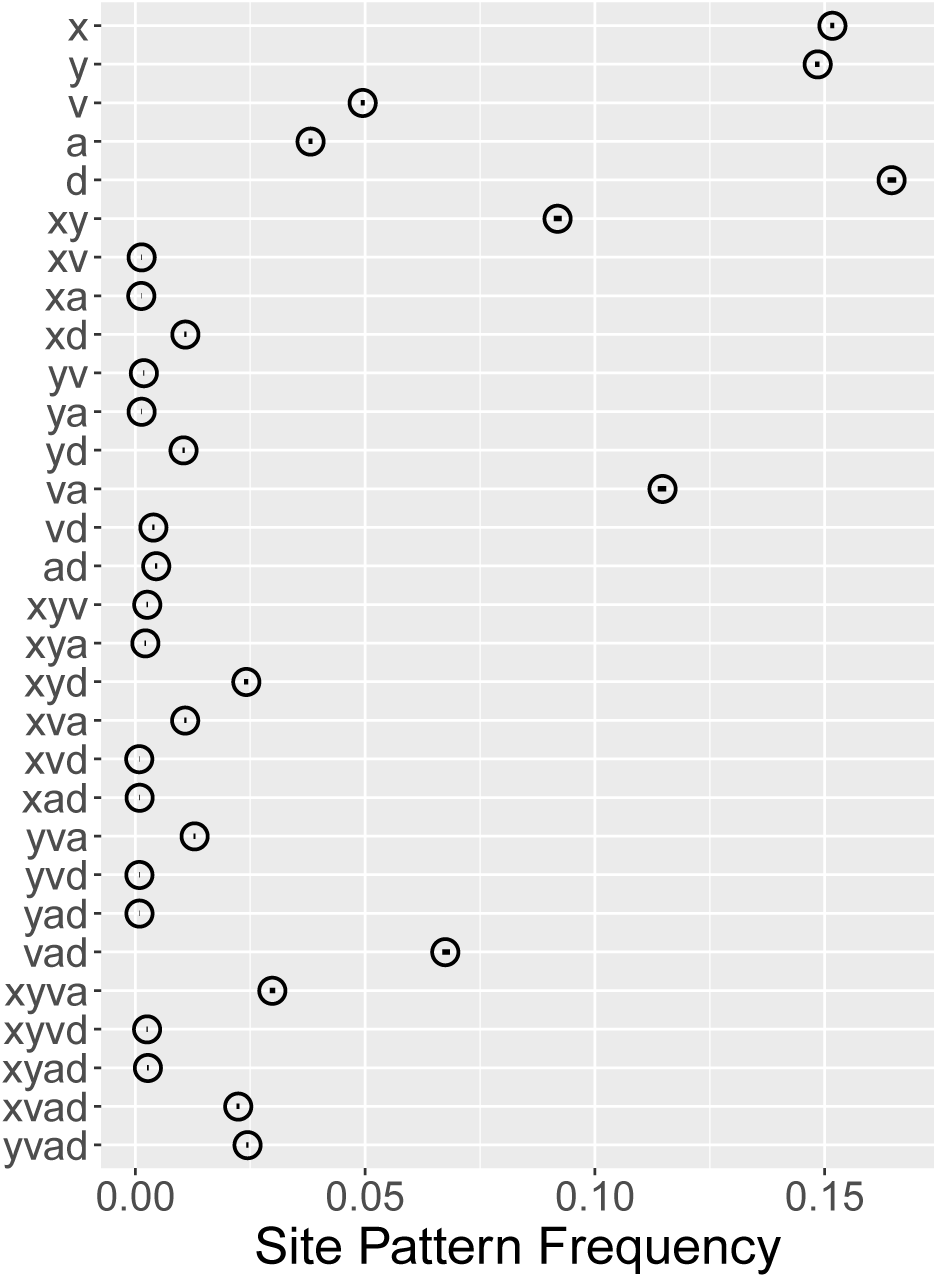
Observed site pattern frequencies. Horizontal axis shows the relative frequency of each site pattern in random samples consisting of a single haploid genome from each of *X, Y, V, A*, and *D*, representing Africa, Europe, Vindija Neanderthal, Altai Neanderthal, Denisovan, and superarchaic. Horizontal lines (which look like dots) are 95% confidence intervals estimated by a moving-blocks bootstrap [11]. Data: Simons Genome Diversity Project [12] and Max Planck Institute for Evolutionary Anthropology [13].

Greek letters in Fig. 1 label episodes of admixture. We label models by concatenating greek letters to indicate the episodes of admixture they include. For example, model “*αβ*” includes only episodes *α* and *β*. Our model does not include gene flow from Denisovans into moderns, because there is little evidence of such gene flow into Europeans [12, 16]. Two years ago we studied a model that included only one episode of admixture: *α*, which refers to gene flow from Neanderthals into Europeans [14]. The left panel of Fig. 3 shows the residuals from this model, using the new data. Several are far from zero, suggesting that something is missing from the model [17].

**Figure 3:**
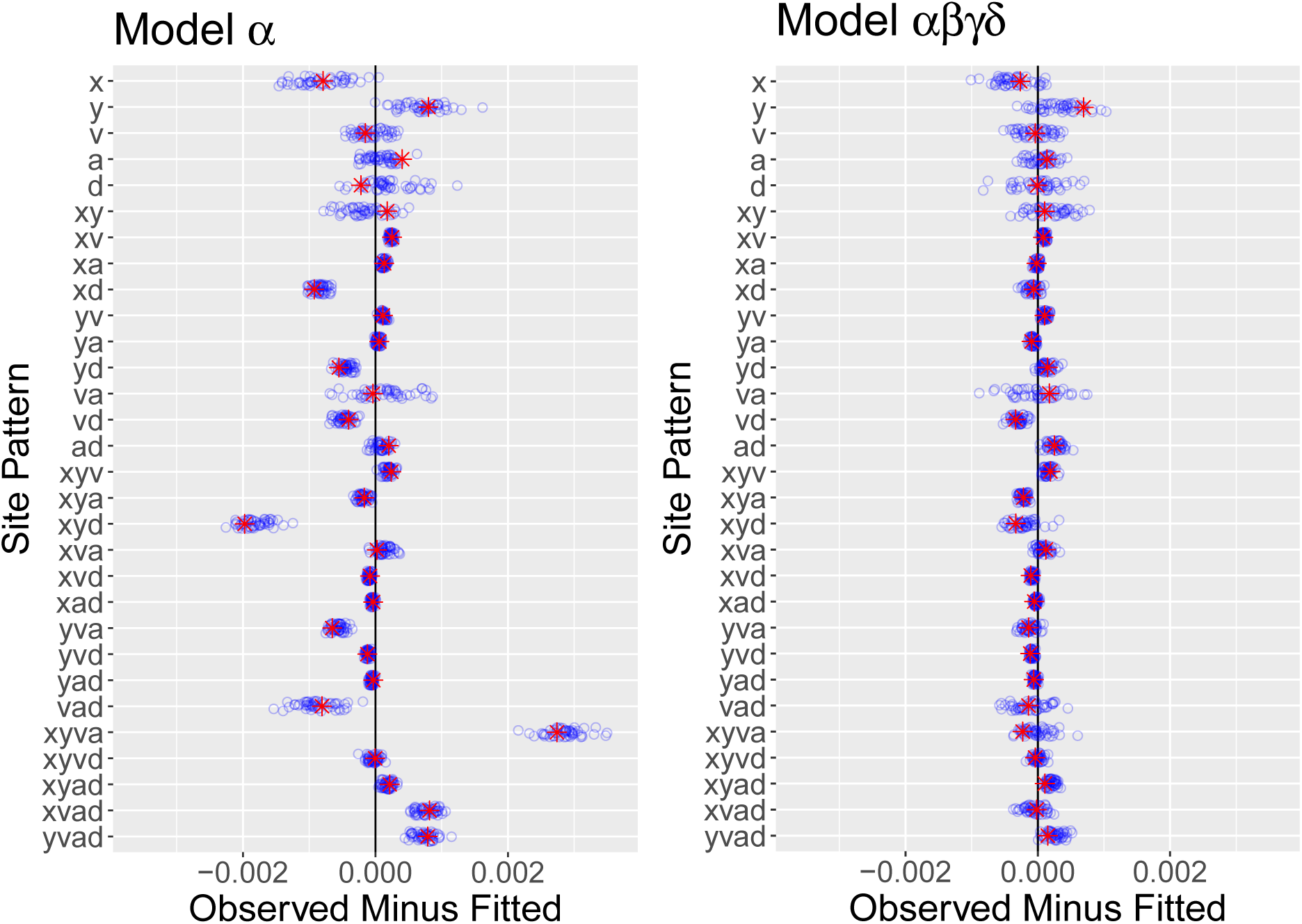
Residuals from models *α* and *αβγδ*. Key: red asterisks, real data; blue circles, 50 bootstrap replicates.

Recent literature suggests some of what might be missing. There is evidence for admixture into Denisovans from a “superarchaic” population, which was distantly related to other humans [18, 19, 2, 13, 20] and also for admixture from early moderns into Neanderthals [20]. These episodes of admixture appear as *β* and *γ* in Fig. 1. Adding *β* and/or *γ* to the model improved the fit, yet none of the resulting models were satisfactory. For example, model *αβγ* implied (implausibly) that superarchaics separated from other hominins seven million years ago.

To understand what might still be missing, consider what we know about the early middle Pleistocene, around 600 kya. At this time, large-brained hominins appear in Europe along with Acheulean stone tools [4, 5]. They were probably African immigrants, because similar fossils and tools occur earlier in Africa. According to one hypothesis, these early Europeans were Neanderthal ancestors [6, 7]. Somewhat earlier—perhaps 750 kya [8, table S12.2]—the “neandersovan” ancestors of Neanderthals and Denisovans separated from the lineage leading to modern humans. Neandersovans may have separated from an African population and then expanded into Eurasia. If so, they would not have been expanding into an empty continent, for Eurasia had been inhabited since 1.85 mya [21]. Neandersovan immigrants may have met the indigenous superarchaic population of Eurasia. This suggests a fourth episode of admixture— from superarchaics into neandersovans—which appears as *δ* in Fig. 1.

## Results

We considered eight models, all of which include *α*, and including all combinations of *β, γ*, and/or *δ*. In choosing among complex models, it is important to avoid overfitting. Conventional methods such as AIC [22] are not available, because we don’t have access to the full likelihood function. Instead, we use the *bootstrap estimate of predictive error* (bepe) [23, 24, 10]. The best model is the one with the lowest value of bepe. When no model is clearly superior, it is better to average across several than to choose just one [25]. For this purpose, we used *bootstrap model averaging* (booma) [25, 10]. The booma weight of the *i*th model is the fraction of data sets (including the real data and 50 bootstrap replicates) in which that model “wins,” i.e. has the lowest value of bepe. The bepe values and booma weights of all models are in table 1.

**Table 1:** Bootstrap estimate of predictive error (bepe) values and bootstrap model average (booma) weights

| Model | bepe | weight |
| --- | --- | --- |
| $\alpha$ | $1.16 \times 10^{-6}$ | 0 |
| $\alpha\delta$ | $0.87 \times 10^{-6}$ | 0 |
| $\alpha\gamma$ | $0.62 \times 10^{-6}$ | 0 |
| $\alpha\gamma\delta$ | $0.44 \times 10^{-6}$ | 0 |
| $\alpha\beta$ | $0.18 \times 10^{-6}$ | 0 |
| $\alpha\beta\gamma$ | $0.17 \times 10^{-6}$ | 0 |
| $\alpha\beta\delta$ | $0.15 \times 10^{-6}$ | 0.16 |
| $\alpha\beta\gamma\delta$ | $0.13 \times 10^{-6}$ | 0.84 |

The best model is *αβγδ*, which includes all four episodes of admixture. It has smaller residuals (Fig. 3, right), the lowest bepe value, and the largest booma weight. One other model— *αβδ*—has a positive booma weight, but all others have weight zero. To understand what this means, recall that bootstrap replicates approximate repeated sampling from the process that generated the data. The models with zero weight lose in *all* replicates, implying that their disadvantage is large compared with variation in repeated sampling. On this basis, we can reject these models. Neither of the two remaining models can be rejected. These results provide strong support for two episodes of admixture (*β* and *δ*) and qualified support for a third (*γ*). Not only does this support previously-reported episodes of gene flow, it also reveals a much older episode, in which neandersovans interbred with superarchaics. Model-averaged parameter estimates, which use the weights in table 1, are graphed in Fig. 4 and listed in Supporting Online Material (SOM) table S1.

**Figure 4:**
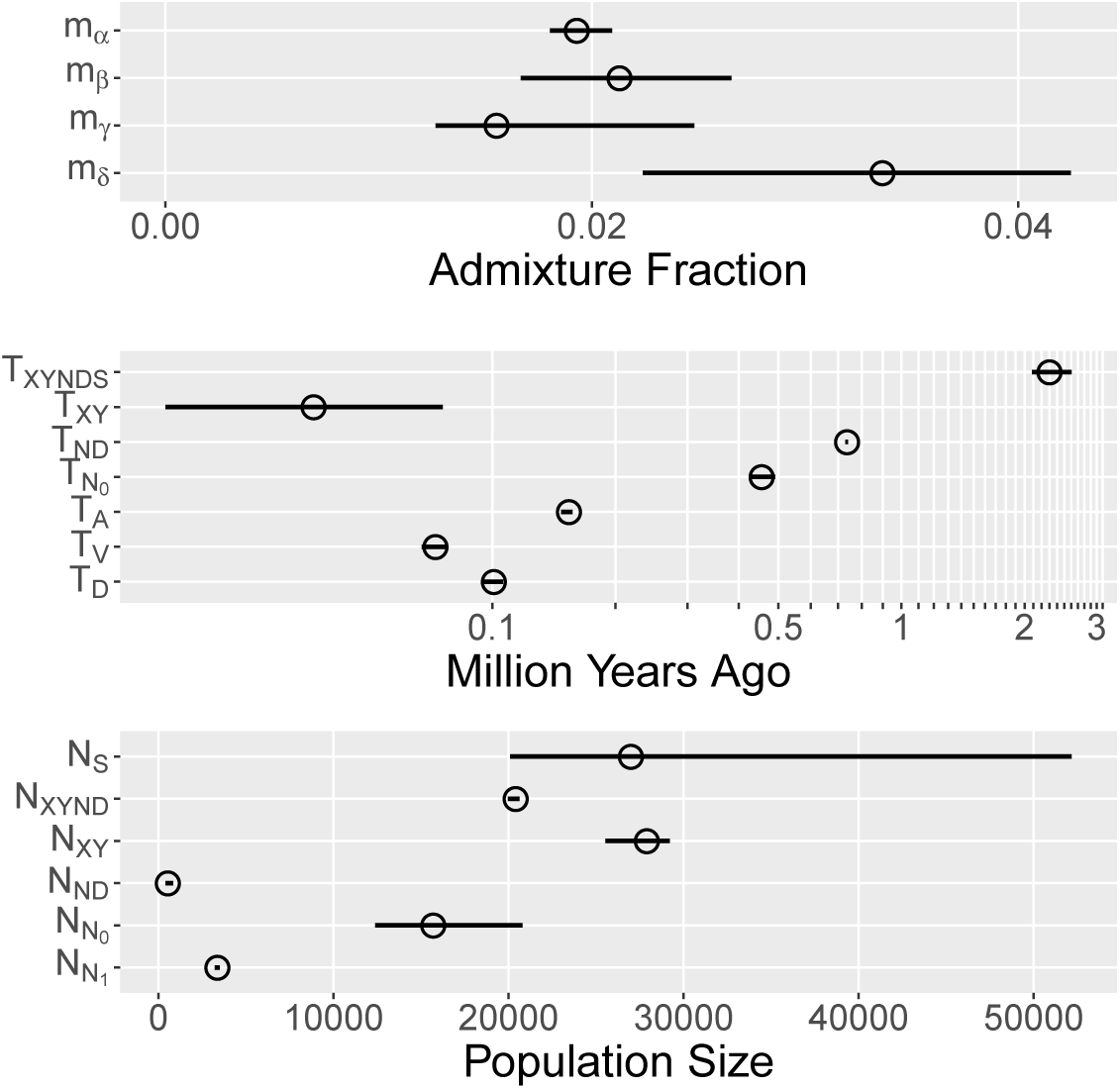
Model-averaged parameter estimates with 95% confidence intervals estimated by moving-blocks bootstrap [11]. Key: *m*_*α*_, fraction of *Y* introgressed from *N*; *m*_*β*_, fraction of *D* introgressed from *S*; *m*_*γ*_, fraction of *N* introgressed from *XY*; *m*_*δ*_; fraction of *ND* introgressed from *S*; *T*_*XY NDS*_, superarchaic separation time; *T*_*XY*_, separation time of *X* and *Y*; *T*_*ND*_, separation time of *N* and *D*; 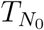, end of early epoch of Neanderthal history; *T*_*A*_, age of Altai Neanderthal fossil; *T*_*V*_, age of Vindija Neanderthal fossil; *T*_*D*_, age of Denisovan fossil; *N*_*S*_, size of superarchaic population; *N*_*XY ND*_, size of populations *XYND* and *XYNDS*; *N*_*XY*_, size of population *XY*; *N*_*ND*_, size of population *ND*; 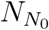, size of early Neanderthal population; 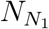, size of late Neanderthal population. Parameters that exist in only one model are not averaged.

Episode *δ*, which proposes gene flow from superarchaics into neandersovans, is a novel hypothesis. Before accepting it, we should ask whether the evidence in its favor could be artifactual, reflecting a bias in site pattern frequencies caused by sequencing error or somatic mutations. Sequencing error adds a positive bias to the frequency of each singleton site pattern proportional to the per-nucleotide error rate in the corresponding population (SOM). Somatic mutations have a similar effect. These biases might explain the evidence for episode *δ*, if it were true that larger values of *m*_*δ*_ (the fraction of superarchaic admixture in neandersovans) imply larger frequencies of singleton site patterns. However, table 2 shows that this is not the case. There is no consistent tendency for singleton frequencies to increase with *m*_*δ*_. Indeed, three of them decrease. Consequently, the evidence that *m*_*δ*_ > 0 cannot be the result of a positive bias in the frequencies of singleton site patterns. The evidence for *δ* admixture cannot be an artifact of sequencing error or somatic mutations.

**Table 2:** Effect on singleton site pattern frequencies of gene flow (*m*_*δ*_) from superarchaics into neandersovans. Column 2 shows expected frequencies of singleton site patterns in a model in which *m*_*δ*_ = 0 and all other parameters are as fitted under model *αβγδ*. In column 3, all parameters including *m*_*δ*_ are as fitted under this model. Column 4 is obtained by subtracting column 2 from column 3. Expected site pattern frequencies were estimated using legosim with 10^7^ iterations.

| Site<br>Pattern | Frequency |  | difference |
| --- | --- | --- | --- |
| | $m_\delta = 0$ | $m_\delta = 0.034$ | |
| x | 0.15583 | 0.15174 | -0.00409 |
| y | 0.15176 | 0.14778 | -0.00398 |
| v | 0.04974 | 0.04942 | -0.00032 |
| a | 0.03795 | 0.03798 | 0.00003 |
| d | 0.16051 | 0.16444 | 0.00393 |

The superarchaic separation time, *T*_*XY NDS*_, has a point estimate of 2.3 mya. This estimate may be biased upward, because our molecular clock assumes a fairly low mutation rate of 0.38×10^−9^ per nucleotide site per year. Other authors prefer slightly higher rates [26]. Although this rate is apparently insensitive to generation time among the great apes, it is sensitive to the age of male puberty. If the average age of puberty during the past two million years were half-way between those of modern humans and chimpanzees, the yearly mutation rate would be close to 0.45 × 10^−9^ [27, Fig. 2B], and our estimate of *T*_*XY NDS*_ would drop to 1.9 mya—just at the origin of the genus *Homo*. Under this clock, the 95% confidence interval is 1.8–2.2 mya.

If superarchaics separated from an African population, then this separation must have preceded the arrival of superarchaics in Eurasia. Nonetheless, our 1.8–2.2 mya interval includes the 1.85 mya date of the earliest Eurasian archaeological remains at Dmanisi [21]. Thus, superarchaics may descend from the earliest human dispersal into Eurasia, as represented by the Dmanisi fossils. On the other hand, some authors prefer a higher mutation rate of 0.5×10^−9^ per year [2]. Under this clock, the lower end of our confidence interval would be 1.6 mya. Thus, our results are also consistent with the view that superarchaics entered Eurasia after the earliest remains at Dmanisi.

Parameter *N*_*S*_ is the effective size of the superarchaic population. This parameter can be estimated because there are two sources of superarchaic DNA in our sample (*β* and *δ*), and this implies that coalescence time within the superarchaic population affects site pattern frequencies. Although this parameter has a broad confidence interval, even the low end implies a fairly large population of about 20,000. This does not require large numbers of superarchaic humans, because effective size can be inflated by geographic population structure [28]. Our large estimate may mean that neandersovans and Denisovans received gene flow from two different superarchaic populations.

Parameter *T*_*ND*_ is the separation time of Neanderthals and Denisovans. Our point estimate— 737 kya—is remarkably old. Furthermore, the neandersovan population that preceded this split was remarkably small: *N*_*ND*_ ≈ 500. This supports our previous results, which indicated an early separation of Neanderthals and Denisovans and a bottleneck among their ancestors [14].

Because our analysis includes two Neanderthal genomes, we can estimate the effective size of the Neanderthal population in two separate epochs. The early epoch extends from 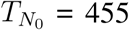 kya to *T*_*ND*_ = 737 kya, and within this epoch the effective size was large: 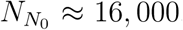. It was smaller during the later epoch: 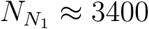. These results support previous findings that the Neanderthal population was large at first but then declined in size [2, 13].

## Discussion

This project began with a puzzle. We had argued in 2017 that Neanderthals and Denisovans separated early, that their neandersovan ancestors endured a bottleneck of population size, and that the post-separation Neanderthal population was large [14]. That analysis omitted singleton site patterns. Mafessoni and Prüfer [15] pointed out that introducing singletons led to different results. In response Rogers et al. [17] agreed, but also observed that the with-singleton analysis implied that the Denisovan fossil was only 4000 years old—a result that is plainly wrong. Furthermore, a residual analysis showed that neither of the models under discussion in 2017 fit the data very well [17]. Something was apparently missing from both models—but what? The present paper provides an answer to that question.

Our results shed light on the early portion of the middle Pleistocene—about 600 kya—when large-brained hominins appear in the fossil record of Europe along with Acheulean stone tools. There is disagreement about how these early Europeans should be interpreted. Some see them as the common ancestors of modern humans and Neanderthals [29], others as an evolutionary dead end, later replaced by immigrants from Africa [30, 31], and others as early representatives of the Neanderthal lineage [6, 7]. Our estimates are most consistent with the last of these views. They imply that by 600 kya Neanderthals were already a distinct lineage, separate not only from the modern lineage but also from Denisovans.

These results resolve a discrepancy involving human fossils from Sima de los Huesos (SH). Those fossils had been dated to at least 350 kya and perhaps 400–500 kya [32]. Genetic evidence showed that they were from a population ancestral to Neanderthals and therefore more recent than the separation of Neanderthals and Denisovans [9]. However, genetic evidence also indicated that this split occurred about 381 kya [2, table S12.2]. This was hard to reconcile with the estimated age of the SH fossils. To make matters worse, improved dating methods later showed that the SH fossils are even older—about 600 ky—and much older than the molecular date of the Neanderthal-Denisovan split [33]. Our estimates resolve this conflict, because they push the date of the split back well beyond the age of the SH fossils.

Our estimate of the Neanderthal-Denisovan separation time conflicts with 381 kya estimate discussed above [2, 15]. This discrepancy results in part from differing calibrations of the molecular clock. Under our clock, the 381 ky date becomes 502 ky [14], but this is still far from our own 737 ky estimate. The remaining discrepancy may reflect differences in our models of history. Misspecified models often generate biased parameter estimates.

Our new results on Neanderthal population size differ from those we published in 2017 [14]. At that time, we argued that the Neanderthal population was substantially larger than others had estimated. Our new estimates are more in line with those published by others [2, 13]. The difference does not result from our new and more elaborate model, because we get similar results from model *α*, which (like our 2017 model) allows only one episode of gene flow (SOM table S2). Instead, it was including the Vindija Neanderthal genome that made the difference. Without this genome, we still get a large estimate 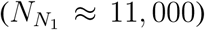, even using model *αβγδ* (SOM table S3). This implies that the Neanderthals who contributed DNA to modern Europeans were more similar to the Vindija Neanderthal than to the Altai Neanderthal, as others have also shown [13].

Our results revise the date at which superarchaics separated from other humans. One previous estimate put this date between 0.9 and 1.4 mya [2, p. 47], which implied that superarchaics arrived well after the initial human dispersal into Eurasia around 1.9 mya. This required a complex series of population movements between Africa and Eurasia [34, pp. 66-71]. Our new estimates do not refute this reconstruction, but they do allow a simpler one, which involves only three expansions of humans from Africa into Eurasia: an expansion of early *Homo* at about 1.9 mya, an expansion of neandersovans at about 700 kya, and an expansion of modern humans at about 50 kya.

Our results indicate that neandersovans interbred with superarchaics early in the middle Pleistocene, shortly after expanding into Eurasia. This is the earliest known admixture between hominin populations. Furthermore, the two populations involved were more distantly related than any pair of human populations previously known to interbreed. According to our estimates, neandersovans and superarchaics had been separate for about 1.2 my. Later, when superarchaics exchanged genes with Denisovans, the two populations had been separate even longer. By comparison, the Neanderthals and Denisovans who interbred with modern humans had been separate less that 0.7 my.

It seems likely that superarchaics descend from the initial human settlement of Eurasia. As discussed above, the large effective size of the superarchaic population hints that it comprised at least two deeply-divided subpopulations, of which one mixed with neandersovans and another with Denisovans. We suggest that around 700 kya, neandersovans expanded from Africa into Eurasia, endured a bottleneck of population size, interbred with indigenous Eurasians, largely replaced them, and separated into eastern and western subpopulations—Denisovans and Neanderthals. These same events unfolded once again around 50 kya as modern humans expanded out of Africa and into Eurasia, largely replacing the Neanderthals and Denisovans.

## Materials and methods

### Study design

Our sample of modern genomes includes Europeans but not other Eurasians. This allows us to avoid modeling gene flow from Denisovans, because there is no evidence of such gene flow into Europeans. The precision of our estimates depends largely on the number of nucleotides studied. For this reason, we use entire high-coverage genomes. The number of genomes sampled per population has little effect on our analyses, because of our focus on the between-population component of genetic variation, i.e. on site-pattern frequencies. Nonetheless, our sample of modern genomes for the Yoruban, French, and English includes all those available from the Simons Genome Diversity Project (SGDP) [12], as detailed in SOM. We also include all available the high-coverage archaic genomes [13]. These data provide extremely accurate estimates of site-pattern frequencies, as indicated by the tiny confidence intervals in Fig. 2. The large confidence intervals for some parameters in Fig. 4 reflect identifiability problems (discussed below) and would not be alleviated by an increase in sample size.

### Quality control (QC)

Our QC pipeline for the SGDP genomes excludes genotypes at which FL equals 0 or N. We also excluded sex chromosomes, normalized all variants at a given nucleotide site using the human reference genome, excluded sites within 7 bases of the nearest INDEL, and included sites only if they are monomorphic or are biallelic SNPs. Further details are provided in SOM. All ancient genomes were also filtered against .bed files, which identify bases that pass the Max Planck QC filters. These .bed files are available at http://ftp.eva.mpg.de/neandertal/Vindija/FilterBed.

### Molecular clock calibration

We assume a mutation rate of 1.1 × 10^−8^ per site per generation [35] and a generation time of 29 y—a yearly rate of 0.38 × 10^−9^. To calibrate the molecular clock, we assume that the modern and neandersovan lineages separated *T*_*XY ND*_ = 25,920 generations before the present [14]. This is based on an average of several PSMC estimates published by Prüfer et al. [2, table S12.2]. The average of their estimates is 570.25 ky, assuming a mutation rate 0.5 × 10^−9^/bp/y. Under our clock, their separation time becomes 751.69 ky or 25,920 generations.

### Statistical analysis

Because of our focus on deep history, we base statistical analyses on site pattern frequencies, using the Legofit statistical package [10]. This method ignores the within-population component of genetic variation and is therefore unaffected by recent changes in population size. For example, the sizes of populations *X, Y*, and *D* (Fig. 1) have no effect, so we need not complicate our model with parameters describing the size histories of these populations. This allows us to focus on the distant past.

Nonetheless, our models are quite complex. For example, model *αβγδ* has 17 free parameters. To choose among models of this complexity, we need methods of residual analysis, model selection, and model averaging. Legofit provides these methods, but alternative methods generally do not. These methods are described in detail elsewhere [10], so we summarize them only briefly here.

We choose among models by minimizing the *bootstrap estimate of predictive error* (bepe) [23, 24]. This approach is needed because we can’t use methods, such as Akaike’s information criterion [22], that depend on likelihood. Bepe is analogous to cross validation, but uses bootstrap replicates instead of partitions of the data. The model is fit to each bootstrap replicate and then tested against the real data, after applying a correction for bootstrap bias. Bepe estimates the mean squared difference between observed and predicted site pattern frequencies, when the model is fit to one data set and tested against another.

We also use *bootstrap model averaging* (booma) [25], which assigns weights to individual models, based on their bepe values. Parameters are estimated as the weighted average of estimates from individual models. The booma weight of the *i*th model is the fraction of replicates (including the real data and 50 bootstrap replicates) in which that model “wins,” i.e. has the lowest value of bepe. Because bootstrap replicates approximate repeated sampling from the process that generated the data, a model will receive weight zero if its disadvantage (as measured by bepe) is large compared with variation in repeated sampling.

SOM Fig. S3 illustrates a problem of statistical identifiability. Several parameters are tightly correlated with others, indicating that our problem has fewer dimensions than parameters. This does not lead to incorrect estimates, but it broadens the confidence intervals of the parameters involved. Legofit addresses this problem by using principal components analysis to remove dimensions that account for less than a fraction 0.001 of the total variance. This narrows confidence intervals and increases the accuracy of parameter estimates.

Uncertainties are estimated by moving blocks bootstrap [11], using a block size of 500 single-nucleotide polymorphisms. Our statistical pipeline is detailed in SOM.

## Supporting information

Supplemental methods and results

## Acknowledgments

We thank Ryan Bohlender, Elizabeth Cashdan, Fabrizio Mafessoni, Nala Rogers, Jon Seger, and Timothy Webster for comments.

## Author contributions

ARR designed the study, did the statistical analyses, and wrote the paper. NSH and AAA developed and employed the quality control pipeline.

## Data and materials availability

All data needed to evaluate the conclusions in the paper are present in the paper, the Supplementary Materials, or at osf.io/vrwna. The Legofit software is available at `github.com:alanrogers/github.git`.

## Human subjects

This project was declared exempt (`IRB_00093972`) from review by the Institutional Review Board of the University of Utah on July 13, 2016.

