## Supplemental methods and results for "Neanderthal-Denisovan ancestors interbred with a distantly-related hominin"

### Methods and materials

#### Samples

All modern genomes are from the Simons Genome Diversity Project (SGDP) [12] and include 3 Yorubans (`SS6004475`, `LP6005442-DNA_B02`, `LP6005442-DNA_A02`), 5 French (`SS6004468`, `LP6005441-DNA_C02`, `LP6005441-DNA_D02`, `LP6005441-DNA_B05`, `LP6005441-DNA_A05`), and 2 English (`LP6005442-DNA_E10`, `LP6005442-DNA_F10`). The ancient genomes in our sample are the Altai Neanderthal genome, the Vindija Neanderthal genome, and the Denisovan genome [13], all of which are available at `http://cdna.eva.mpg.de/neandertal/Vindija/VCF`.

#### Outgroup

We used the chimpanzee [36] and gorilla [37, 38] reference genomes to call ancestral alleles. These genomes, aligned to human hg19, are available at the Santa Cruz Genome Browser. We downloaded .axt files aligned to human autosomes and generated a single .raf file for the chimpanzee and gorilla genomes. The chimpanzee pipeline is on p. 6, and the gorilla pipeline was similar. We then generated a combined chimpanzee-gorilla .raf file as follows:

```
joinraf chimp.raf.gz gorilla.raf.gz | gzip -c > chigor.raf
```

In this pipeline, `axt2raf.py` and `joinraf` are from the Legofit package [10].

#### Quality Control (QC)

QC is of special concern in this project because we include singleton site patterns, whose counts may be inflated by sequencing error. Sequencing errors tend to show up as heterozygous sites in diploid genomes. To explore this effect, we graphed heterozygosity against a measure of genotype quality.

The SGDP data files measure genotype quality in terms of the "FL value." Fig. S1 shows that heterozygosity is inflated only for FL values 0 and N. Our QC pipeline for the SGDP therefore excludes these values.

The slurm script on p. 8 was used to filter the Altai Neanderthal genome and convert it to .raf format. The QC pipelines for the Vindija Neanderthal and Denisovan genomes were similar.

The QC pipeline for SGDP data is illustrated on p. 9. We generated a separate .raf file for each genome. We then used `joinraf`, from the Legofit package [10] to combine these into two files: `frengland.raf.gz` is a combined .raf file for the French and English samples, and `yoruba.raf.gz` is for the Yoruban samples.

#### Site pattern frequencies

The program `sitepat`, a component of the Legofit package [10], was used to call ancestral alleles, tabulate site patterns, and generate bootstrap replicates (see p. 11). The resulting site-pattern frequencies are shown in Fig. 2 of the main text. These data underlie all the analyses described here.

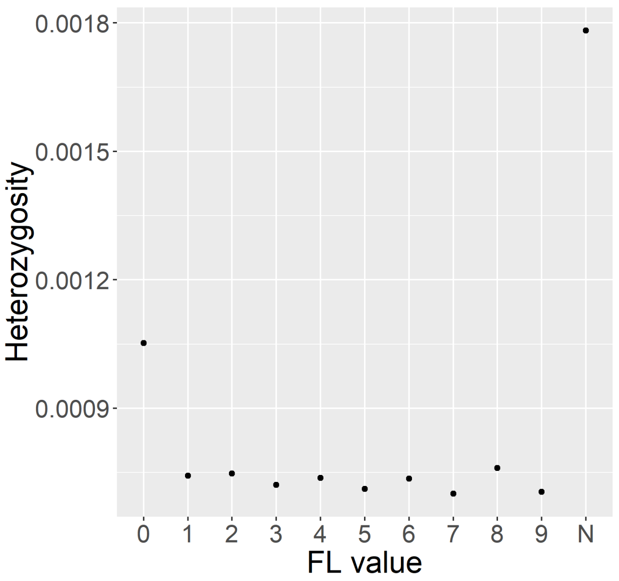


Figure S1: **Heterozygosity as a function of FL value for genome SS6004468 of the SGDP [12].**

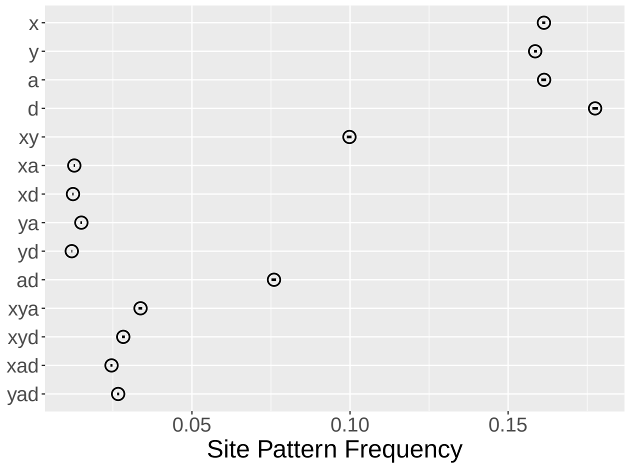


Figure S2: **Observed site pattern frequencies excluding the Vindija genome.** For details, see Fig. 2 of the main text.

Those data may seem counterintuitive, because they include two Neanderthal genomes rather than just one. For this reason, mutations shared exclusively by Europeans and Neanderthals are split between site patterns $ya$, $yv$, and $yav$, so each of these patterns has a lower frequency than the corresponding pattern would have in an analysis involving only one Neanderthal. The same is not true of Denisovans, because we have only one Denisovan genome. Consequently, $yd$ is more common than $yv$, and $yd$ than $ya$, in spite of the well-documented evidence for gene flow into Europeans from Neanderthals but not Denisovans. The data in Fig. S2 are more intuitively accessible, because there is only one Neanderthal genome. Note that in these data $ya$ is more common than $yd$, implying that Europeans share more derived alleles with Neanderthals than with Denisovans. Also, $xd$ and $xa$ are about equally frequent (implying that Africans are equally related to the two archaics), and $ya$ is more common than $xa$ (implying that Europeans share more with Neanderthals than Africans do).

#### Legofit analyses

To distinguish between the Legofit package [10] and the legofit program within that package, we capitalize the former but not the latter. The legofit program was used to estimate parameters. The process began by using a text editor to construct a .lgo file, which describes the history of population size, subdivision, and admixture. The file for model $\alpha\beta\gamma\delta$ is on p. 12. The analysis proceeded in four stages, which were parallelized on a cluster computer. The overall estimation pipeline is a shell script, pipeline.sh (p. 13), which launches a series of slurm scripts on the cluster.

In each stage of the analysis, a separate legofit job was run for each of 51 data sets: the real data and 50 bootstrap replicates. The first two stages identified associations among parameters (illustrated in Fig. S3). These associations imply that the maximization problem has fewer dimensions than parameters, so after stage 2 we re-expressed all free variables in terms of a reduced set of principal components. In each legofit run, composite likelihood was maximized using the *differential evolution* (DE) [39] algorithm. This algorithm maintains a swarm of points, each of which represents a point in parameter space.

Stage 1 of the analysis began with points in the DE swarm scattered widely across parameter space. The objective function was evaluated with only modest precision. It is possible that some of these legofit jobs converged onto different local maxima of the composite likelihood surface. At the end of stage 1, each legofit job wrote its own swarm of points to a file. This stage was done by two slurm scripts: a1.slr (p. 14) and a1boot.slr (p. 14).

In stage 2, each legofit job initialized its DE swarm by reading all the state files produced during stage 1. This allowed stage 2 to choose among the local optima discovered during stage 1. During stage 2, each evaluation of the objective function was done to high precision. The slurm scripts for stage 2 are: a2.slr (p. 15) and a2boot.slr (p. 16).

After stage 2, `pclgo` was used to re-express all free variables in terms of principal components, excluding components that account for less than a fraction 0.001 of the total variance. The slurm script for this step is pclgo.slr (p. 18). It generated file b.lgo (p. 18).

Stages 3 and 4 were analogous to stages 1 and 2. Stage 3 operated at modest precision and generated 51 state files. Then stage 4, which operated at high precision, read these state files so that legofit could choose among local optima. The slurm scripts for stages 3 and 4 are: b1.slr (p. 21), b1boot.slr (p. 22), b2.slr (p. 22), and b2boot.slr (p. 24).

After all four stages had run for a particular model, bepe was run as follows:

```
LC_ALL=C
bepe ../xyvad.opf ../boot/boot*.opf -L b2.legofit b2boot*.legofit > b2.bepe
```

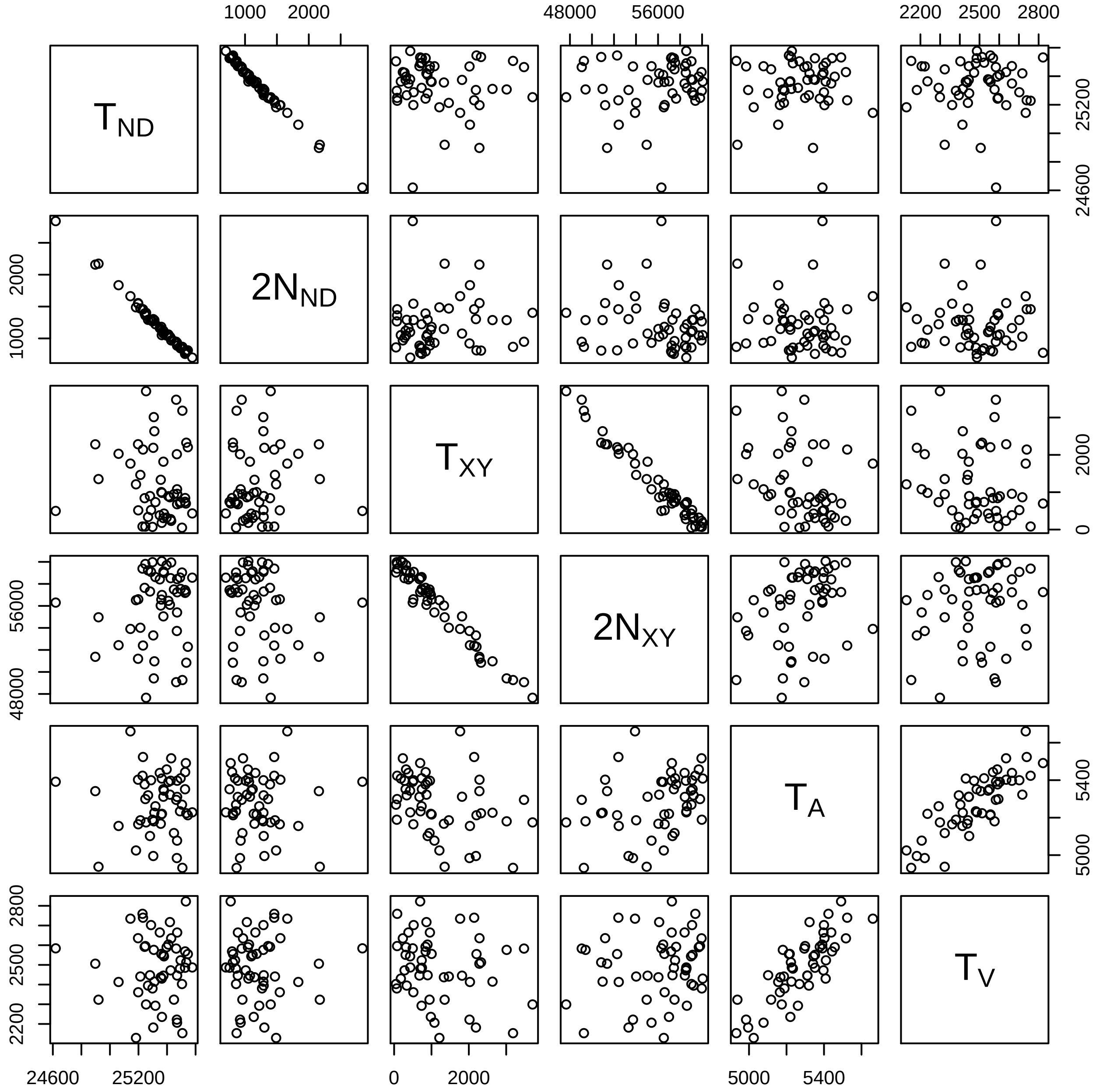


Figure S3: **Associations between estimates of several pairs of parameters after second stage in analysis of model $\alpha\beta\gamma\delta$.** Parameters are defined in Fig. 4 of the main text.

Setting `LC_ALL=C` ensured that the bash shell would sort input files in a consistent order on different machines. This produced file b2.bepe. The first line in this file gives the bepe value for the real data, and the remaining lines give bepe values for bootstrap replicates. The bepe values in table 1 of the main text were taken from the top lines of the b2.bepe files for the various models.

For each model, we also created a file called b2.flat, which contains estimates for each parameter (in columns) and each data set (in rows). This was done as follows:

```
LC_ALL=C
flatfile.py b2.legofit b2boot*.legofit > b2.flat
```

On our server, we keep the analysis of each model in its own directory. These directories have names such as `a` (for model $\alpha$), `ab` (for $\alpha\beta$), and so on. From the parent directory, we executed booma as follows:

```
booma */b2.bepe -F */b2.flat > all.bma
```

This created file all.bma, which provided the booma weights in table 1 of the main text.

Finally, we used the Legofit program bootci.py:

```
bootci.py all.bma > all.bootci
```

This generated all.bootci, which contains model-averaged point estimates and confidence intervals for all parameters.

#### Effect of sequencing error and somatic mutations

Sequencing errors and somatic mutations have similar effects, since both tend to occur as rare variants that appear only once in the sample. This section asks how these processes affect site pattern frequencies. We focus on sequencing error, with the understanding that similar comments apply to somatic mutation.

Although all site patterns may be affected by sequencing error, the effect is largest on singleton site patterns. Because the vast majority of the genome is monomorphic, most errors occur at monomorphic sites. These errors usually appear as single, spurious heterozygotes at sites that are otherwise monomorphic. Such sites contribute only to singleton site patterns. Let us evaluate the magnitude of this effect.

Consider the singleton site pattern $x$, which refers to the case in which the derived allele is present in a random nucleotide sampled from population $X$ but is absent in homologous nucleotides sampled from the other populations. This site pattern occurs at site $j$ with probability $z_{xj} = p_{Xj} q_{Yj} q_{Vj} q_{Aj} q_{Dj}$, where $p_{Xj}$ is the frequency of the derived allele at site $j$ among samples from $X$, and the $q$s are homologous ancestral allele frequencies among samples from Europe, the Vindija Neanderthal, the Altai Neanderthal, and Denisovan [10]. To tabulate site pattern frequencies, we sum $z_{xj}$ across the genome, and then normalize so that the frequencies sum to unity.

At the vast majority of sites, all sampled nucleotides are copies of the ancestral allele, so that (absent sequencing error) $p_{Xj} = 0$, $q_{Yj} = q_{Vj} = q_{Aj} = q_{Dj} = 1$, and $z_{xj} = 0$. Suppose however that a sequencing error occurs at site $j$ in one of the $k_X$ genomes sampled from population $X$, generating a spurious heterozygote. In this case, $p_{Xj} = z_{xj} = 1/2k_X$. Such a site would contribute $1/2k_X$ to site pattern $x$ but would contribute nothing to other site patterns.

How frequent are such errors across the genome? Let $v_X^{(i)}$ denote the probability that a homozygous site is incorrectly called as a heterozygote in the $i$th genome sampled from population $X$. With approximate probability $\sum_{i=1}^{k_X} v_X^{(i)}$, one of the $k_X$ genomes from population $X$ is a spurious

Table S1: Model-averaged parameter estimates. Time is measured in units of generations. Parameters are defined in Fig. 4 of the main text.

| Parameter | Estimate | 95% Confidence Interval | |
|---|---|---|---|
| | | Low | High |
| $m_\alpha$ | 0.0193 | 0.0180 | 0.0210 |
| $m_\beta$ | 0.0213 | 0.0167 | 0.0265 |
| $m_\gamma$ | 0.0155 | 0.0127 | 0.0248 |
| $m_\delta$ | 0.0336 | 0.0224 | 0.0425 |
| $T_A$ | 5310.3627 | 5078.2072 | 5414.5073 |
| $T_V$ | 2505.6239 | 2317.3653 | 2655.7058 |
| $T_D$ | 3480.3598 | 3242.3082 | 3665.4764 |
| $T_{N_0}$ | 15706.0431 | 14593.5069 | 16956.3343 |
| $T_{\text{ND}}$ | 25400.0216 | 25135.4015 | 25526.0436 |
| $T_{\text{XY}}$ | 1262.1696 | 548.0009 | 2610.7864 |
| $T_{\text{XYNDS}}$ | 79334.5216 | 71901.2093 | 89764.7578 |
| $2N_{N_0}$ | 31411.1823 | 24766.9196 | 41586.1098 |
| $2N_{N_1}$ | 6744.8324 | 6444.4014 | 6994.2331 |
| $2N_{ND}$ | 1081.0078 | 796.8532 | 1680.0852 |
| $2N_S$ | 53994.6216 | 40180.9289 | 104336.0927 |
| $2N_{XY}$ | 55818.3059 | 51085.2402 | 58415.9833 |
| $2N_{\text{XYND}}$ | 40830.8412 | 39916.6608 | 41264.0956 |

heterozygote at some given nucleotide site. (This assumes $v_X^{(i)}$ is small enough that we can ignore higher-order terms.) The expected contribution of sequencing error to $z_{xj}$ is the product of $1/2k_X$ and $\sum_{i=1}^{k_X} v_X^{(i)}$, which equals $\bar{v}_X/2$, where $\bar{v}_X = \frac{1}{k_X}\sum_{i=1}^{k_X} v_X^{(i)}$ is the mean error rate in population $X$.

In summary, sequencing errors cause a positive bias in $z_{xj}$, which is on average equal to $\bar{v}_X/2$. This generates a positive bias in estimates of singleton site pattern frequencies. The bias in site pattern frequencies is somewhat larger, because the process of normalizing frequencies involves multiplying by a factor greater than unity. Nonetheless, it remains true that the frequencies of singleton site patterns have a positive bias, which is proportional to the per-nucleotide error rate. This error rate may vary among populations, so some singleton site patterns may have a larger bias than others. Somatic mutations would add a similar bias.

### Supplementary results

Table S1 shows model-averaged parameter estimates and confidence intervals. These values are graphed in Fig. 4 of the main text. Table S2 shows parameter estimates under model $\alpha$. Table S3 shows estimates under model $\alpha\beta\gamma\delta$ using a data set that excludes the Vindija Neanderthal genome.

### Chimpanzee genome

```
#SBATCH --time=24:00:00
#SBATCH --nodes 1
#SBATCH --ntasks 1
#SBATCH -o chimp.raf.gz
```

Table S2: Estimates under model $\alpha$.

| Parameter | Estimate | 95% Confidence Interval Low | High |
|---|---|---|---|
| $m_\alpha$ | 0.0233 | 0.0214 | 0.0247 |
| $T_A$ | 5832.6400 | 5611.7700 | 5924.1675 |
| $T_V$ | 2901.1200 | 2822.2825 | 3152.0650 |
| $T_D$ | 2547.8300 | 2391.5175 | 2866.2300 |
| $T_{N_0}$ | 11872.3000 | 11874.9750 | 14477.2250 |
| $T_{\text{ND}}$ | 25830.4000 | 25626.8500 | 25844.0000 |
| $T_{\text{XY}}$ | 9119.8000 | 7968.9250 | 9627.7225 |
| $2N_{N_0}$ | 19008.9000 | 18851.6250 | 24242.1000 |
| $2N_{N_1}$ | 5764.8800 | 5756.7425 | 6636.8075 |
| $2N_{ND}$ | 240.9980 | 205.0327 | 780.7515 |
| $2N_{XY}$ | 31509.6000 | 30403.5750 | 34034.3500 |
| $2N_{\text{XYND}}$ | 42702.1000 | 41913.7250 | 42963.9750 |

Table S3: Estimates under model $\alpha\beta\gamma\delta$ with a data set that excludes the Vindija Neanderthal genome.

| Parameter | Estimate | 95% Confidence Interval Low | High |
|---|---|---|---|
| $m_\alpha$ | 0.0280 | 0.0216 | 0.0300 |
| $m_\beta$ | 0.0121 | 0.0078 | 0.0191 |
| $m_\gamma$ | 0.0370 | 0.0159 | 0.0597 |
| $m_\delta$ | 0.0150 | 0.0108 | 0.0224 |
| $T_A$ | 5212.1500 | 4978.5525 | 5250.3600 |
| $T_D$ | 3559.0700 | 3305.9675 | 3608.0500 |
| $T_{\text{ND}}$ | 25096.8000 | 24691.3000 | 25449.7000 |
| $T_{\text{XY}}$ | 2676.8100 | 1362.5525 | 3616.3100 |
| $T_{\text{XYNDS}}$ | 99076.5000 | 79243.9250 | 114293.2500 |
| $2N_N$ | 22214.4000 | 13151.2500 | 26884.1750 |
| $2N_{ND}$ | 1749.4000 | 980.6828 | 2674.1275 |
| $2N_S$ | 30590.0000 | 2699.3275 | 29870.5500 |
| $2N_{XY}$ | 49007.9000 | 46390.1000 | 53327.5000 |
| $2N_{\text{XYND}}$ | 40245.1000 | 39554.3000 | 40910.1500 |

```
#SBATCH -e chimpraf.err

# ensure consistent sort
export LC_ALL=C

# format of axt files for reference genome.
fmt="panTro5/chr%d.hg19.panTro5.net.axt.gz\n"

# concatenate axt files
seq 22 | sort | xargs printf $fmt | xargs zcat |

# make .raf.gz file
axt2raf.py |

# get rid of "chr" in names of chromosomes
sed 's/^chr//' |

# compress
gzip -c
```

#### QC pipeline for the Altai Neanderthal genome

```
#!/bin/bash
#SBATCH -J amkraf
#SBATCH --account=rogersa-kp
#SBATCH --partition=rogersa-kp
#SBATCH --time=36:00:00
#SBATCH --nodes 1
#SBATCH --ntasks 1
#SBATCH -o altai.raf.gz
#SBATCH -e mkraf.err

# create scratch directory
scrdir=/scratch/general/lustre/$USER/$SLURM_JOBID
mkdir -p $scrdir

# reference genome directory
refdir=$HOME/group/rogers/data/hg19

# copy ref to scratch and uncompress; also copy index file
ref_fa=${scrdir}/hg19-sort.fa
twoBitToFa ${refdir}/hg19-sort.2bit $ref_fa
cp ${refdir}/hg19-sort.fa.fai $scrdir

vfmt="chr%d_mq25_mapab100.vcf.gz\n"

# The new bed file names begin with "chr". The old ones start with
```

```
# integers.
bfmt="../bed/chr%d_mask.bed.gz\n"

# ensure consistent sort
export LC_ALL=C

# concatenate vcf files
seq 22 | sort | xargs printf ${vfmt} | xargs bcftools concat |

# keep only sites listed in bed files
bedtools intersect -header -sorted -a "stdin" \
-b <(seq 22 | sort | xargs printf ${bfmt} | xargs zcat) |

# Combine lines that refer to different alleles at a single position
bcftools norm --multiallelics +any --fasta-ref $ref_fa -Ou |

# We only want only SNPs and REFs not adjacent to indels
# This genome doesn't have INFO/MQ. File name suggests MQ>=25
bcftools filter --SnpGap 7 --set-GTs '.' \
    --include '(TYPE="snp"||TYPE="ref") & FMT/GQ>29' -Ou |

# print in format required by raf
bcftools query -f '%CHROM\t%POS\t%REF\t%ALT[\t%GT]\n' |

# generate raf .raf.gz file
raf | gzip -c

# delete scratch files
rm -rf $scrdir
```

#### QC pipeline for modern genomes from the SGDP

```
#!/bin/bash
#SBATCH -J yri
#SBATCH --account=rogersa
#SBATCH --partition=kingspeak
#SBATCH --time=48:00:00
#SBATCH --nodes 1
#SBATCH --ntasks 2
#SBATCH -o yri%a.out # stdout
#SBATCH -e yri%a.err # stderr

i=${SLURM_ARRAY_TASK_ID}

# Set and make a temporary directory in the scratch
SCRDIR=/scratch/general/lustre/$USER/$SLURM_JOB_ID
mkdir -p $SCRDIR
```

```
# get current directory
loc=`pwd`

# reference genome with chromosomes sorted in lexical order
cp ~/harris/hg19-sort.fa $SCRDIR

# Retrieve the names of the samples.
# yri.txt is a file that contains, in its first column, the IDs
# of the genomes to be included in this data set.
peeps=(`cut yri.txt -f1`)

# The array ID for slurm is used to subset the samples to a single
# individual
peep=${peeps["$i"]}
ref=hg19-sort.fa

# Copy the sample to the scratch
target="$peep".sorted.vcf.gz
cp $target $SCRDIR

# Print name of the target file to out and err files so we don't have
# to keep track of which slurm ID corresponds to which sample
echo "$target"
>&2 echo "$target"

# Switch to the scratch drive for speed
cd $SCRDIR

#combine records of different types into a single line, then they will
#be removed by the filter command alone. Indels are normaliszed to the
#same positions using the refernence.
bcftools norm --multiallelics +any --fasta-ref $ref $target |

# Remove low quality and missing data.
bcftools filter -e 'FMT/FL = "0" | FMT/FL = "N"' |

# Limit the data to SNPs and fixed sites.
bcftools filter -i 'TYPE="snp" | TYPE="ref"' --SnpGap 7 |

# Output the columns that are required by raf
bcftools query -f '%CHROM\t%POS\t%REF \t%ALT\t[%GT]\n' |

# Make raf file
raf | gzip > "$peep".raf.gz

# Copy the resulting file to the original directory
cp "$peep".raf.gz "$loc"/"$peep".raf.gz
```

```
# Remove the temporary scratch directory.
rm -rf $SCRDIR
```

#### Calling ancestral alleles, tabulating site patterns, and generating bootstrap replicates

```
#!/bin/bash
#SBATCH -J sitepat
#SBATCH --account=rogersa-kp
#SBATCH --partition=rogersa-kp
#SBATCH --time=36:00:00
#SBATCH --nodes 1
#SBATCH --ntasks 1
#SBATCH -o xyvad.opf # stdout
#SBATCH -e sitepat.err # stderr

# create scratch directory
#scrdir=/scratch/general/lustre/$USER/$SLURM_JOBID
scrdir=/scratch/general/lustre/$USER
mkdir -p $scrdir

# uncompress raf files
gunzip -c $HOME/group/harris/data/yri/yoruba.raf.gz > $scrdir/yoruba.raf
gunzip -c $HOME/group/harris/data/eur/frengland.raf.gz > $scrdir/frengland.raf
gunzip -c $HOME/group/rogers/data/vindija/orig/vindija.raf.gz > $scrdir/vindija.raf
gunzip -c $HOME/group/rogers/data/altai/orig2/altai.raf.gz > $scrdir/altai.raf
gunzip -c $HOME/group/rogers/data/denisova/orig2/denisova.raf.gz > $scrdir/denisova.raf
gunzip -c $HOME/group/rogers/data/ape/chigor.raf.gz > $scrdir/chigor.raf

echo "finished decompression" 1>&2

# create observed pattern frequency (.opf) file and bootstrap replicates
mkdir -p boot
time sitepat -1 --bootfile boot/boot --bootreps 50 \
x=$scrdir/yoruba.raf \
y=$scrdir/frengland.raf \
v=$scrdir/vindija.raf \
a=$scrdir/altai.raf \
d=$scrdir/denisova.raf \
    outgroup=$scrdir/chigor.raf

echo "finished sitepat" 1>&2

# delete scratch files
#rm -rf $scrdir
```

### Estimation

#### Input file for model $\alpha\beta\gamma\delta$: a.lgo

This file is used in estimation stages 1 and 2.

```
# Gene flow: N->Y, S->D, XY->N, S->ND
# S = superarchaic; XY = early modern; ND = Neandersovan; Y = Europe;
# A, Altai; V=Vindija; D=Denisovan. Altai and Vindija on same branch
# TmN < Tv < Ta
time fixed  zero = 0
twoN fixed  one = 1
time fixed  TmN = 1        # no coalesc. events can happen b/t 0 and Tv
time fixed  Txynd = 25920  # \citet[table~S12.2, p.~90]{Li:N-505-43-S88}
time free   Txynds = 60000 # arbitrary: 1.74 myr
time free   Tnd = 24000
time free   Tav = 14000
time free   Txy = 10000
time free   Td  = 2500
time free   Ta  = 4000
time free   Tv  = 1000
twoN free   twoNav = 2000
twoN free   twoNn = 2000
twoN free   twoNnd = 2000
twoN free   twoNxy = 20000
twoN free   twoNxynd = 20000
twoN free   twoNs = 2000
mixFrac free   mN = 0.01
mixFrac free   mS = 0.01
mixFrac free   mSND = 0.01
mixFrac free   mXY = 0.01
time constrained TmXY = 0.5*(Txy + Tav)
time constrained TmS = 0.5*(Td + Tnd)
time constrained TmSND = 0.5*(Tnd + Txynd)
segment x    t=zero   twoN=one     samples=1
segment y    t=zero   twoN=one     samples=1
segment v    t=Tv     twoN=twoNn   samples=1
segment a    t=Ta     twoN=twoNn   samples=1
segment a2   t=TmXY   twoN=twoNn
segment d    t=Td     twoN=one     samples=1
segment d2   t=TmS    twoN=one
segment y2   t=TmN    twoN=one
segment n    t=TmN    twoN=twoNn
segment s    t=TmS    twoN=twoNs
segment s2   t=TmSND  twoN=twoNs
segment av   t=Tav    twoN=twoNav
segment nd   t=Tnd    twoN=twoNnd
segment nd2  t=TmSND  twoN=twoNnd
segment xy   t=Txy    twoN=twoNxy
```

```
segment xy2  t=TmXY   twoN=twoNxy
segment xynd t=Txynd  twoN=twoNxynd
segment xynds t=Txynds twoN=twoNxynd
mix    d    from d2 + mS * s
mix    y    from y2 + mN * n
mix    a    from a2 + mXY * xy2
mix    nd   from nd2 + mSND * s2
derive n    from v
derive v    from a
derive a2   from av
derive av   from nd
derive d2   from nd
derive x    from xy
derive y2   from xy
derive xy   from xy2
derive xy2  from xynd
derive nd2  from xynd
derive s    from s2
derive s2   from xynds
derive xynd from xynds
```

##### Overall pipeline

```
#!/bin/bash

# Slurm pipeline.  sbatch returns a string of form "Submitted batch
# job 123456". We only want the last word in this string. The sbatch
# steps below initially set a variable equal to the entire
# string. Then we extract the last word with a command like j1=${j1##*
# }. The syntax here is obscure to me but seems to work.

export SBATCH_ACCOUNT=rogersa-kp
export SBATCH_PARTITION=rogersa-kp

# Stage 1 of analysis.
j1=$(sbatch a1.slr)
j2=$(sbatch --array=0-49 a1boot.slr)
j1=${j1##* }
j2=${j2##* }

# Stage 2 of analysis.
j3=$(sbatch --dependency=afterok:$j1:$j2 a2.slr)
j4=$(sbatch --dependency=afterok:$j1:$j2 --array=0-49 a2boot.slr)
j3=${j3##* }
j4=${j4##* }

# Rewrite .lgo file in terms of principal components. This makes
# file b.lgo.
```

```
j5=$(sbatch --dependency=afterok:$j3:$j4 pclgo.slr)
j5=${j5##* }

# Stage 1 for model b
j6=$(sbatch --dependency=afterok:$j5 b1.slr)
j7=$(sbatch --dependency=afterok:$j5 --array=0-49 b1boot.slr)
j6=${j6##* }
j7=${j7##* }

# Stage 2 for model b
sbatch --dependency=afterok:$j6:$j7 b2.slr
sbatch --dependency=afterok:$j6:$j7 --array=0-49 b2boot.slr
```

##### Stage 1 legofit job: a1.slr

```
#!/bin/bash
#SBATCH -J ABCDa1
#SBATCH --account=rogersa-kp
#SBATCH --partition=rogersa-kp
#SBATCH --time=36:00:00
#SBATCH --nodes 1
#SBATCH --ntasks 1
#SBATCH -o a1.legofit # stdout
#SBATCH -e a1.err # stderr

ifile=../xyvad.opf
stateout=a1.state
lgofile=a.lgo

time legofit -1 --stateOut $stateout --tol 3e-5 \
  -S 5000@10000 -S 100@100000 -S 1000@2000000 $lgofile $ifile
```

##### Stage 1 legofit for bootstrap replicates: a1boot.slr

```
#!/bin/bash
#SBATCH -J ABCDa1boot
#SBATCH --account=rogersa-kp
#SBATCH --partition=rogersa-kp
#SBATCH --time=36:00:00
#SBATCH --nodes 1
#SBATCH --ntasks 1
#SBATCH -o a1boot%a.legofit # stdout
#SBATCH -e a1boot%a.err # stderr

i=${SLURM_ARRAY_TASK_ID}
ifile=$(printf "../boot/boot%d.opf" $i)    # input file
stateout=$(printf "a1boot%d.state" $i)
lgofile=a.lgo
```

```
time legofit -1 --stateOut $stateout --tol 3e-5 \
  -S 5000@10000 -S 100@100000 -S 1000@2000000 $lgofile $ifile
```

##### Stage 2 legofit job: a2.slr

```
#!/bin/bash
#SBATCH -J ABCDa2
#SBATCH --account=rogersa-kp
#SBATCH --partition=rogersa-kp
#SBATCH --time=36:00:00
#SBATCH --nodes 1
#SBATCH --ntasks 1
#SBATCH -o a2.legofit # stdout
#SBATCH -e a2.err # stderr

ifile=../xyvad.opf
lgofile=a.lgo

time legofit -1 --tol 2e-5 -S 1000@2000000 $lgofile $ifile \
  --stateIn a1.state \
  --stateIn a1boot0.state \
  --stateIn a1boot1.state \
  --stateIn a1boot10.state \
  --stateIn a1boot11.state \
  --stateIn a1boot12.state \
  --stateIn a1boot13.state \
  --stateIn a1boot14.state \
  --stateIn a1boot15.state \
  --stateIn a1boot16.state \
  --stateIn a1boot17.state \
  --stateIn a1boot18.state \
  --stateIn a1boot19.state \
  --stateIn a1boot2.state \
  --stateIn a1boot20.state \
  --stateIn a1boot21.state \
  --stateIn a1boot22.state \
  --stateIn a1boot23.state \
  --stateIn a1boot24.state \
  --stateIn a1boot25.state \
  --stateIn a1boot26.state \
  --stateIn a1boot27.state \
  --stateIn a1boot28.state \
  --stateIn a1boot29.state \
  --stateIn a1boot3.state \
  --stateIn a1boot30.state \
  --stateIn a1boot31.state \
  --stateIn a1boot32.state \
```

```
  --stateIn a1boot33.state \
  --stateIn a1boot34.state \
  --stateIn a1boot35.state \
  --stateIn a1boot36.state \
  --stateIn a1boot37.state \
  --stateIn a1boot38.state \
  --stateIn a1boot39.state \
  --stateIn a1boot4.state \
  --stateIn a1boot40.state \
  --stateIn a1boot41.state \
  --stateIn a1boot42.state \
  --stateIn a1boot43.state \
  --stateIn a1boot44.state \
  --stateIn a1boot45.state \
  --stateIn a1boot46.state \
  --stateIn a1boot47.state \
  --stateIn a1boot48.state \
  --stateIn a1boot49.state \
  --stateIn a1boot5.state \
  --stateIn a1boot6.state \
  --stateIn a1boot7.state \
  --stateIn a1boot8.state \
  --stateIn a1boot9.state
```

##### Stage 2 legofit for bootstrap replicates: a2boot.slr

```
#!/bin/bash
#SBATCH -J ABCDa2boot
#SBATCH --account=rogersa-kp
#SBATCH --partition=rogersa-kp
#SBATCH --time=36:00:00
#SBATCH --nodes 1
#SBATCH --ntasks 1
#SBATCH -o a2boot%a.legofit # stdout
#SBATCH -e a2boot%a.err # stderr

i=${SLURM_ARRAY_TASK_ID}
ifile=`printf "../boot/boot%d.opf" $i`    # input file
lgofile=a.lgo

time legofit -1 --tol 2e-5 -S 1000@2000000 $lgofile $ifile \
  --stateIn a1.state \
  --stateIn a1boot0.state \
  --stateIn a1boot1.state \
  --stateIn a1boot10.state \
  --stateIn a1boot11.state \
  --stateIn a1boot12.state \
  --stateIn a1boot13.state \
```

```
--stateIn a1boot14.state \
--stateIn a1boot15.state \
--stateIn a1boot16.state \
--stateIn a1boot17.state \
--stateIn a1boot18.state \
--stateIn a1boot19.state \
--stateIn a1boot2.state \
--stateIn a1boot20.state \
--stateIn a1boot21.state \
--stateIn a1boot22.state \
--stateIn a1boot23.state \
--stateIn a1boot24.state \
--stateIn a1boot25.state \
--stateIn a1boot26.state \
--stateIn a1boot27.state \
--stateIn a1boot28.state \
--stateIn a1boot29.state \
--stateIn a1boot3.state \
--stateIn a1boot30.state \
--stateIn a1boot31.state \
--stateIn a1boot32.state \
--stateIn a1boot33.state \
--stateIn a1boot34.state \
--stateIn a1boot35.state \
--stateIn a1boot36.state \
--stateIn a1boot37.state \
--stateIn a1boot38.state \
--stateIn a1boot39.state \
--stateIn a1boot4.state \
--stateIn a1boot40.state \
--stateIn a1boot41.state \
--stateIn a1boot42.state \
--stateIn a1boot43.state \
--stateIn a1boot44.state \
--stateIn a1boot45.state \
--stateIn a1boot46.state \
--stateIn a1boot47.state \
--stateIn a1boot48.state \
--stateIn a1boot49.state \
--stateIn a1boot5.state \
--stateIn a1boot6.state \
--stateIn a1boot7.state \
--stateIn a1boot8.state \
--stateIn a1boot9.state
```

##### Use pclgo to rewrite .lgo file: pclgo.slr

This slurm script generates file b.lgo, which re-expresses all free variables in terms of principal components.

```
#!/bin/bash
#SBATCH -J ABCDpclgo
#SBATCH --account=rogersa-kp
#SBATCH --partition=rogersa-kp
#SBATCH --time=00:10:00
#SBATCH --nodes 1
#SBATCH --ntasks 1
#SBATCH -e pclgo.err # stderr

# Make b.lgo. Exclude dimensions that account for less than
# a fraction 0.001 of the variance.
cmd="(grep ^# a.lgo;"
cmd+=" pclgo --tol 0.001 a.lgo a2.legofit a2boot*.legofit;"
cmd+=" grep -v ^# a.lgo | egrep -v \"\<free\>\")"
echo "# "$cmd > b.lgo
eval $cmd >> b.lgo
```

##### The model re-expressed in terms of principal components: b.lgo

```
# (grep ^# a.lgo; pclgo --tol 0.001 a.lgo a2.legofit a2boot*.legofit; grep -v ^# a.lgo | egrep
# Gene flow: N->Y, S->D, XY->N, S->ND
# S = superarchaic; XY = early modern; ND = Neandersovan; Y = Europe;
# A, Altai; V=Vindija; D=Denisovan. Altai and Vindija on same branch
# TmN < Tv < Ta
# PCA calculated with gsl_linalg_SV_decomp
# Fraction of variance:
#      pc1      pc2      pc3      pc4      pc5      pc6      pc7      pc8
#  0.28617  0.16142  0.13504  0.09936  0.08719  0.07037  0.04574  0.03928
#      pc9     pc10     pc11     pc12     pc13     pc14     pc15     pc16
#  0.02687  0.01799  0.01319  0.00697  0.00585  0.00306  0.00101  0.00044
#     pc17
#  0.00004
param free [    -4,      4] pc1 = 0
param free [    -4,      4] pc2 = 0
param free [    -4,      4] pc3 = 0
param free [    -2,      2] pc4 = 0
param free [    -2,      2] pc5 = 0
param free [    -2,      2] pc6 = 0
param free [    -2,      2] pc7 = 0
param free [    -2,      2] pc8 = 0
param free [    -2,      2] pc9 = 0
param free [    -2,      2] pc10 = 0
param free [    -2,      2] pc11 = 0
param free [    -1,      1] pc12 = 0
```

```
param free [    -1,      1] pc13 = 0
param free [    -1,      1] pc14 = 0
param free [    -1,      1] pc15 = 0
time constrained Txynds = 80931.461 + 3215.4589*pc1 + 161.65839*pc2 +
    1046.5493*pc3 - 2065.5586*pc4 - 434.78469*pc5 - 653.6697*pc6 +
    1468.7086*pc7 + 1673.0571*pc8 + 1415.4685*pc9 + 160.79802*pc10 +
    1974.9601*pc11 - 4217.3768*pc12 + 688.08865*pc13 + 4554.1119*pc14 -
    1814.0015*pc15
time constrained Tnd = 25334.194 - 33.790649*pc1 + 51.27568*pc2 -
    47.252127*pc3 - 80.974156*pc4 + 34.768373*pc5 + 1.9789971*pc6 +
    42.798412*pc7 + 5.4043439*pc8 + 23.79307*pc9 - 12.884031*pc10 -
    13.059141*pc11 + 6.054159*pc12 + 1.733805*pc13 - 10.896501*pc14 +
    1.1266543*pc15
time constrained Tav = 16585.857 - 18.024609*pc1 - 456.06418*pc2 -
    28.692127*pc3 - 209.38598*pc4 + 102.36818*pc5 + 26.676295*pc6 -
    20.395005*pc7 - 94.841348*pc8 + 1.2264252*pc9 + 25.143582*pc10 -
    90.774545*pc11 + 7.5990123*pc12 + 183.67019*pc13 - 210.20911*pc14 -
    560.90489*pc15
time constrained Txy = 1191.4813 + 265.07115*pc1 + 136.98478*pc2 -
    238.28159*pc3 + 242.23856*pc4 + 402.61829*pc5 - 79.649622*pc6 -
    8.4959931*pc7 - 149.98977*pc8 - 15.581845*pc9 + 101.73441*pc10 -
    163.43127*pc11 + 100.22147*pc12 - 3.3787396*pc13 + 208.71263*pc14 -
    40.478798*pc15
time constrained Td = 3496.8824 - 29.550471*pc1 - 5.5383938*pc2 +
    41.497169*pc3 + 6.4285721*pc4 + 34.26318*pc5 - 85.344952*pc6 -
    33.826473*pc7 - 30.279575*pc8 + 103.13788*pc9 - 21.897014*pc10 +
    26.446686*pc11 + 10.30263*pc12 - 20.155073*pc13 - 3.1067375*pc14 -
    1.0735719*pc15
time constrained Ta = 5279.6584 - 39.224542*pc1 - 16.323958*pc2 +
    59.535082*pc3 + 7.8138125*pc4 + 59.025697*pc5 + 13.006729*pc6 +
    27.800519*pc7 - 15.953376*pc8 - 29.772804*pc9 - 45.774969*pc10 -
    1.5863634*pc11 - 6.1230856*pc12 + 95.048205*pc13 + 29.878247*pc14 +
    39.704553*pc15
time constrained Tv = 2486.7135 - 24.511249*pc1 - 9.6972927*pc2 +
    70.117489*pc3 + 0.63371286*pc4 + 74.019405*pc5 + 15.633188*pc6 +
    44.463088*pc7 + 33.959807*pc8 - 37.399669*pc9 + 29.329041*pc10 +
    7.441393*pc11 - 10.145682*pc12 - 99.18374*pc13 - 9.5116455*pc14 -
    17.027713*pc15
twoN constrained twoNav = 32497.847 + 1379.2188*pc1 - 1460.3469*pc2 +
    39.752646*pc3 - 2331.8037*pc4 - 92.455928*pc5 + 652.77903*pc6 +
    178.81721*pc7 - 4550.9175*pc8 - 949.8518*pc9 - 172.18009*pc10 +
    1211.7284*pc11 + 581.61798*pc12 - 1666.6067*pc13 + 932.71666*pc14 +
    1649.5183*pc15
twoN constrained twoNn = 6943.3176 - 26.387832*pc1 - 75.891896*pc2 -
    39.869486*pc3 - 13.420159*pc4 + 23.747533*pc5 - 7.4974051*pc6 -
    1.1500775*pc7 + 37.256213*pc8 + 28.20307*pc9 + 110.50624*pc10 +
    4.5482975*pc11 + 13.974404*pc12 + 21.066474*pc13 + 32.836787*pc14 +
    63.980599*pc15
```

```
twoN constrained twoNnd = 1210.7456 + 85.223627*pc1 - 110.252*pc2 +
    109.3222*pc3 + 173.54084*pc4 - 70.336708*pc5 - 4.6334948*pc6 -
    87.865666*pc7 - 13.581821*pc8 - 47.387662*pc9 + 34.415643*pc10 +
    35.490281*pc11 - 16.678026*pc12 - 0.66692864*pc13 +
    7.5358424*pc14 + 0.20139938*pc15
twoN constrained twoNxy = 55949.939 - 1089.2405*pc1 - 431.96148*pc2 +
    555.73716*pc3 - 739.4499*pc4 - 1386.7936*pc5 + 212.65038*pc6 +
    34.860967*pc7 + 465.21476*pc8 + 94.382166*pc9 - 243.56779*pc10 +
    575.83637*pc11 - 332.11094*pc12 - 207.79554*pc13 - 249.67258*pc14 +
    633.81552*pc15
twoN constrained twoNxynd = 40481.841 - 110.20278*pc1 -
    7.8870509*pc2 - 202.64252*pc3 + 139.11032*pc4 + 67.762449*pc5 +
    9.0219043*pc6 + 48.355745*pc7 - 39.120808*pc8 - 58.698653*pc9 -
    32.009776*pc10 + 356.19198*pc11 - 96.839309*pc12 + 9.2639439*pc13 -
    52.933393*pc14 - 52.787159*pc15
twoN constrained twoNs = 52104.703 + 742.45756*pc1 - 1368.6566*pc2 -
    1695.4322*pc3 - 2877.2064*pc4 - 3567.2827*pc5 - 24170.851*pc6 +
    6019.7872*pc7 + 1405.666*pc8 - 14841.032*pc9 + 439.81963*pc10 -
    1455.3417*pc11 + 303.05508*pc12 + 992.21143*pc13 - 1817.5284*pc14 +
    1210.5895*pc15
mixFrac constrained mN = 0.01937248 - 8.0947378e-05*pc1 +
    0.00029020798*pc2 + 9.9439299e-05*pc3 - 0.00035000327*pc4 +
    0.00022809347*pc5 - 1.836772e-05*pc6 - 0.00084071313*pc7 +
    7.5594134e-05*pc8 - 0.00034811049*pc9 + 0.00018793504*pc10 +
    0.00018066577*pc11 - 0.00010058954*pc12 + 9.2863e-05*pc13 +
    2.4516309e-05*pc14 - 4.8506313e-05*pc15
mixFrac constrained mS = 0.0200033 - 0.0016630107*pc1 +
    0.0002204065*pc2 - 0.00036922537*pc3 + 0.00066184686*pc4 -
    0.00032500251*pc5 - 0.00012962814*pc6 - 0.0001789951*pc7 -
    0.0016339999*pc8 - 5.2738462e-05*pc9 + 0.0007921886*pc10 -
    0.0017611125*pc11 - 0.0028883776*pc12 - 0.00036314185*pc13 +
    0.00025334317*pc14 - 9.6381029e-05*pc15
mixFrac constrained mSND = 0.033958602 - 0.0040600701*pc1 -
    0.0010118566*pc2 - 0.0013570687*pc3 + 0.00068500249*pc4 -
    0.0014024539*pc5 + 2.0210482e-05*pc6 - 0.0011148578*pc7 +
    0.00045749112*pc8 - 0.0012901906*pc9 - 0.0021240869*pc10 -
    0.00069886122*pc11 + 0.0033960257*pc12 - 0.0019684206*pc13 +
    0.0067152524*pc14 - 0.0027408498*pc15
mixFrac constrained mXY = 0.018767467 + 0.00054508118*pc1 -
    0.0014978088*pc2 - 0.0011236462*pc3 - 0.00015700189*pc4 +
    0.00085292527*pc5 - 2.3646903e-05*pc6 - 0.00093622921*pc7 +
    0.0011169819*pc8 - 7.8259029e-05*pc9 - 0.0021188324*pc10 -
    0.00069880493*pc11 - 0.0010785383*pc12 - 0.00089580326*pc13 -
    0.00020400452*pc14 + 0.0010857181*pc15
time fixed  zero = 0
twoN fixed  one = 1
time fixed  TmN = 1        # no coalesc. events can happen b/t 0 and Tv
time fixed  Txynd = 25920  # \citet[table~S12.2, p.~90]{Li:N-505-43-S88}
```

```
time constrained TmXY = 0.5*(Txy + Tav)
time constrained TmS = 0.5*(Td + Tnd)
time constrained TmSND = 0.5*(Tnd + Txynd)
segment x    t=zero   twoN=one     samples=1
segment y    t=zero   twoN=one     samples=1
segment v    t=Tv     twoN=twoNn   samples=1
segment a    t=Ta     twoN=twoNn   samples=1
segment a2   t=TmXY   twoN=twoNn
segment d    t=Td     twoN=one     samples=1
segment d2   t=TmS    twoN=one
segment y2   t=TmN    twoN=one
segment n    t=TmN    twoN=twoNn
segment s    t=TmS    twoN=twoNs
segment s2   t=TmSND  twoN=twoNs
segment av   t=Tav    twoN=twoNav
segment nd   t=Tnd    twoN=twoNnd
segment nd2  t=TmSND  twoN=twoNnd
segment xy   t=Txy    twoN=twoNxy
segment xy2  t=TmXY   twoN=twoNxy
segment xynd t=Txynd  twoN=twoNxynd
segment xynds t=Txynds twoN=twoNxynd
mix    d    from d2 + mS * s
mix    y    from y2 + mN * n
mix    a    from a2 + mXY * xy2
mix    nd   from nd2 + mSND * s2
derive n    from v
derive v    from a
derive a2   from av
derive av   from nd
derive d2   from nd
derive x    from xy
derive y2   from xy
derive xy   from xy2
derive xy2  from xynd
derive nd2  from xynd
derive s    from s2
derive s2   from xynds
derive xynd from xynds
```

##### Stage 3 legofit job: b1.slr

```
#!/bin/bash
#SBATCH -J ABCDb1
#SBATCH --account=rogersa-kp
#SBATCH --partition=rogersa-kp
#SBATCH --time=36:00:00
#SBATCH --nodes 1
#SBATCH --ntasks 1
```

```
#SBATCH -o b1.legofit # stdout
#SBATCH -e b1.err # stderr

ifile=../xyvad.opf
stateout=b1.state
lgofile=b.lgo

time legofit -1 --stateOut $stateout --tol 3e-5 \
  -S 5000@10000 -S 100@100000 -S 1000@2000000 $lgofile $ifile
```

##### Stage 3 legofit for bootstrap replicates: b1boot.slr

```
#!/bin/bash
#SBATCH -J ABCDb1boot
#SBATCH --account=rogersa-kp
#SBATCH --partition=rogersa-kp
#SBATCH --time=36:00:00
#SBATCH --nodes 1
#SBATCH --ntasks 1
#SBATCH -o b1boot%a.legofit # stdout
#SBATCH -e b1boot%a.err # stderr

i=${SLURM_ARRAY_TASK_ID}
ifile=$(printf "../boot/boot%d.opf" $i)    # input file
stateout=$(printf "b1boot%d.state" $i)
lgofile=b.lgo

time legofit -1 --stateOut $stateout --tol 3e-5 \
  -S 5000@10000 -S 100@100000 -S 1000@2000000 $lgofile $ifile
```

##### Stage 4 legofit job: b2.slr

```
#!/bin/bash
#SBATCH -J ABCDb2
#SBATCH --account=rogersa-kp
#SBATCH --partition=rogersa-kp
#SBATCH --time=36:00:00
#SBATCH --nodes 1
#SBATCH --ntasks 1
#SBATCH -o b2.legofit # stdout
#SBATCH -e b2.err # stderr

ifile=../xyvad.opf
lgofile=b.lgo

time legofit -1 --tol 2e-5 -S 1000@2000000 $lgofile $ifile \
  --stateIn b1.state \
  --stateIn b1boot0.state \
```

```
--stateIn b1boot1.state \
--stateIn b1boot10.state \
--stateIn b1boot11.state \
--stateIn b1boot12.state \
--stateIn b1boot13.state \
--stateIn b1boot14.state \
--stateIn b1boot15.state \
--stateIn b1boot16.state \
--stateIn b1boot17.state \
--stateIn b1boot18.state \
--stateIn b1boot19.state \
--stateIn b1boot2.state \
--stateIn b1boot20.state \
--stateIn b1boot21.state \
--stateIn b1boot22.state \
--stateIn b1boot23.state \
--stateIn b1boot24.state \
--stateIn b1boot25.state \
--stateIn b1boot26.state \
--stateIn b1boot27.state \
--stateIn b1boot28.state \
--stateIn b1boot29.state \
--stateIn b1boot3.state \
--stateIn b1boot30.state \
--stateIn b1boot31.state \
--stateIn b1boot32.state \
--stateIn b1boot33.state \
--stateIn b1boot34.state \
--stateIn b1boot35.state \
--stateIn b1boot36.state \
--stateIn b1boot37.state \
--stateIn b1boot38.state \
--stateIn b1boot39.state \
--stateIn b1boot4.state \
--stateIn b1boot40.state \
--stateIn b1boot41.state \
--stateIn b1boot42.state \
--stateIn b1boot43.state \
--stateIn b1boot44.state \
--stateIn b1boot45.state \
--stateIn b1boot46.state \
--stateIn b1boot47.state \
--stateIn b1boot48.state \
--stateIn b1boot49.state \
--stateIn b1boot5.state \
--stateIn b1boot6.state \
--stateIn b1boot7.state \
--stateIn b1boot8.state \
```

```
  --stateIn b1boot9.state
```

##### Stage 4 legofit for bootstrap replicates: b2boot.slr

```
#!/bin/bash
#SBATCH -J ABCDb2boot
#SBATCH --account=rogersa-kp
#SBATCH --partition=rogersa-kp
#SBATCH --time=36:00:00
#SBATCH --nodes 1
#SBATCH --ntasks 1
#SBATCH -o b2boot%a.legofit # stdout
#SBATCH -e b2boot%a.err # stderr

i=${SLURM_ARRAY_TASK_ID}
ifile=`printf "../boot/boot%d.opf" $i`    # input file
lgofile=b.lgo

time legofit -1 --tol 2e-5 -S 1000@2000000 $lgofile $ifile \
  --stateIn b1.state \
  --stateIn b1boot0.state \
  --stateIn b1boot1.state \
  --stateIn b1boot10.state \
  --stateIn b1boot11.state \
  --stateIn b1boot12.state \
  --stateIn b1boot13.state \
  --stateIn b1boot14.state \
  --stateIn b1boot15.state \
  --stateIn b1boot16.state \
  --stateIn b1boot17.state \
  --stateIn b1boot18.state \
  --stateIn b1boot19.state \
  --stateIn b1boot2.state \
  --stateIn b1boot20.state \
  --stateIn b1boot21.state \
  --stateIn b1boot22.state \
  --stateIn b1boot23.state \
  --stateIn b1boot24.state \
  --stateIn b1boot25.state \
  --stateIn b1boot26.state \
  --stateIn b1boot27.state \
  --stateIn b1boot28.state \
  --stateIn b1boot29.state \
  --stateIn b1boot3.state \
  --stateIn b1boot30.state \
  --stateIn b1boot31.state \
  --stateIn b1boot32.state \
  --stateIn b1boot33.state \
```

```
--stateIn b1boot34.state \
--stateIn b1boot35.state \
--stateIn b1boot36.state \
--stateIn b1boot37.state \
--stateIn b1boot38.state \
--stateIn b1boot39.state \
--stateIn b1boot4.state \
--stateIn b1boot40.state \
--stateIn b1boot41.state \
--stateIn b1boot42.state \
--stateIn b1boot43.state \
--stateIn b1boot44.state \
--stateIn b1boot45.state \
--stateIn b1boot46.state \
--stateIn b1boot47.state \
--stateIn b1boot48.state \
--stateIn b1boot49.state \
--stateIn b1boot5.state \
--stateIn b1boot6.state \
--stateIn b1boot7.state \
--stateIn b1boot8.state \
--stateIn b1boot9.state
```
